# Delta/Theta band EEG activity shapes the rhythmic perceptual sampling of auditory scenes

**DOI:** 10.1101/2020.07.21.213694

**Authors:** Cora Kubetschek, Christoph Kayser

## Abstract

Many studies speak in favor of a rhythmic mode of listening, by which the encoding of acoustic information is structured by rhythmic neural processes at the time scale of about 1 to 4 Hz. Indeed, psychophysical data suggest that humans sample acoustic information in extended soundscapes not uniformly, but weigh the evidence at different moments for their perceptual decision at the time scale of about 2 Hz. We here test the critical prediction that such rhythmic perceptual sampling is directly related to the state of ongoing brain activity prior to the stimulus. Human participants judged the direction of frequency sweeps in 1.2 s long soundscapes while their EEG was recorded. Computing the perceptual weights attributed to different epochs within these soundscapes contingent on the phase or power of pre-stimulus oscillatory EEG activity revealed a direct link between the 4Hz EEG phase and power prior to the stimulus and the phase of the rhythmic component of these perceptual weights. Hence, the temporal pattern by which the acoustic information is sampled over time for behavior is directly related to pre-stimulus brain activity in the delta/theta band. These results close a gap in the mechanistic picture linking ongoing delta band activity with their role in shaping the segmentation and perceptual influence of subsequent acoustic information.

## Introduction

Perception and cognition are controlled by rhythmic activity in the brain^1–3^. These rhythmic processes can reflect directly in behavioral data, such as periodic changes in reaction times or measures of perceptual accuracy relative to stimulus onset^4–7^. More frequently, they are revealed by systematic relations between signatures of rhythmic brain activity and measures of performance, such as changes in accuracy or sensitivity with the power or timing of pre-stimulus activity^8–10^. Concerning hearing, several studies have shown that performance varies with pre-stimulus activity below 10 Hz. For example, participants’ ability to detect brief acoustic targets or to discriminate two subsequent tones varied with the power and phase of brain activity below about 4 Hz^8,9,11–14^. The apparent match between the time scales of perceptual sensitivity and those at which neural activity shapes hearing^15,16^ is seen as strong support of a rhythmic mode of hearing. Such a rhythmic mode could facilitate the amplification of specific (e.g. expected) stimuli and mediate the alignment of endogenous neural activity to the regularities of structured sounds such as speech^17,18^.

A critical prediction based on these studies, and motivated by a link between rhythmic network activity and the functional gain of individual neurons, is that perception should sample acoustic information rhythmically rather than continuously over time^2,10,17,19,20^. Thereby, also information in longer soundscapes that are devoid of an explicit temporal structure should be weighted at precisely those timescales at which rhythmic brain activity is predictive of behavior (i.e. between the delta and theta bands between about 1 and 4 Hz). Studies on speech have provided evidence in favor of this hypothesis^18,21–24^, e.g. by showing that delta band activity serves the chunking or segmentation of speech on a sentence-level time scale^15,18,25^ while theta band activity reflects the processing of syllable-scale information. However, the underlying processes may possibly be specific to speech, which is intrinsically predictive on multiple time scales. Other studies have used periodic sounds to entrain rhythmic neural processes and have shown the persistent and periodic influence of these on behavior for several cycles even after the offset of the entraining sound^26,27^. However, this does not demonstrate a direct influence of spontaneous brain activity on a subsequent rhythmic mode of listening.

To more broadly address the question of whether listening samples acoustic information based on rhythmic processes in the delta or theta time scales, we have previously designed a paradigm allowing the quantification of the moment-by-moment influence of acoustic evidence on perceptual judgments^28^. In that earlier study, we found evidence in favor of a rhythmic listening mode in human participants. However, by design this study did not link the rhythmic weighting of acoustic evidence to brain activity and made the strong assumption that the temporal weighting profile is idiosyncratic across trials^28^. That is, it assumed that the relative sampling phase is consistent on a trial by trial basis. However, if the excitability of auditory pathways is controlled by (rhythmic) pre-stimulus brain activity^29,30^, this assumption could be violated: the temporal perceptual weighting profile at which the momentary acoustic evidence is sampled should change on a trial by trial basis relative to the trial-wise pattern of pre-stimulus activity.

Here we directly tested this prediction by asking whether the rhythmic behavioral use of acoustic information is directly related to pre-stimulus activity. To probe this, we combined psychophysical reverse correlation with EEG recordings obtained while human participants judged the direction of frequency sweeps in pseudo-random soundscapes of 1.2 s duration. We first reproduced our previous results providing evidence for a rhythmic perceptual sampling of extended soundscapes at a frequency of about 2 Hz. Then, we show that the relative timing of these perceptual weights is significantly shaped by the power and phase of pre-stimulus EEG activity at the same time scale, with the perceptual weights of opposing phase bins differing by about 90 degrees.

## Methods

### Participants

The experiment combined a previously described behavioral task with electroencephalography (EEG) recordings in 20 participants (11 females; 19-32 years old). The study was conducted in accordance with the Declaration of Helsinki and was approved by the ethics committee of Bielefeld University. Data was collected with the participants’ written informed consent. Participants reported normal hearing and received monetary compensation of 10 Euro/hour. During the experiment they sat in an electrically and acoustically shielded room (Ebox, Desone, Germany).

### Stimuli

The stimuli and task have been described in detail before^28^. The stimuli were presented via headphones (Sennheiser HD200 Pro) at an average intensity of 65 dB root mean square level. Each stimulus was composed of 30 simultaneously presented sequences of four tones each, whereby each sequence either increased or decreased in frequency over four steps **(Figure 1A**). Each tone had a duration of 30 ms. The starting frequency of each sequence was drawn independently between 128 Hz and 16,384 Hz and increased or decreased in steps of 20 cents. The starting position within each sequence (1 to 4) of the initial 30 sequences were selected at random to ensure that the frequencies, and sequence start/end, were independent across the 30 sequences. Also, the exact starting times of individual tones within a sequence were varied up to 30 ms. To create the impression of an overall frequency sweep over time, the proportion of in/decreasing sequences was systematically varied. This fraction was coded between zero and one, with zero indicating that all sequences decreased and a value of one indicating that all sequences increased in frequency. Each trial was characterized by the actual direction of change (increasing or decreasing) and the associated evidence, coded as the deviation from an ambiguous stimulus (0.5). The design allowed us to vary the amount of sensory evidence between and within a trial around each participant’s threshold (see below). Specifically, each soundscape of 1200 ms duration was divided into ten ‘epochs’, each lasting 120 ms (see **Fig. 1B**). The motion evidence for each epoch was drawn randomly and independently from a Gaussian distribution centered around the participants’ threshold and an SD of 0.2. To obtain the stimulus for a given trial we first determined the sweep direction (increasing or decreasing) and then sampled the motion evidence for each epoch and subsequently generated the sequences of pure tones to fit those parameters.

**Figure 1.**
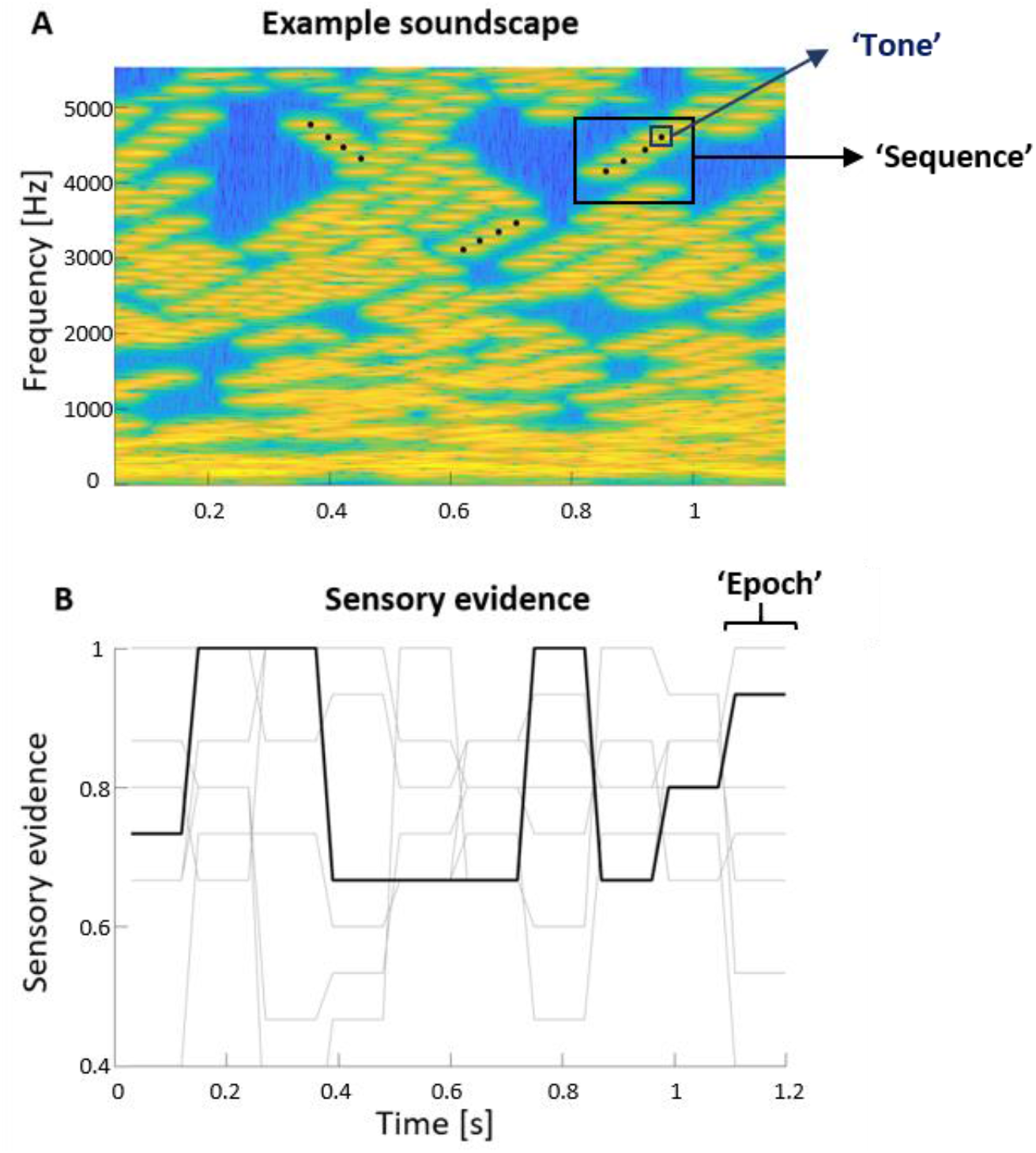
Example soundscape. **A**) Time-frequency representation of one soundscape with an overall increasing frequency sweep. Black dots indicate the four consecutive tones of three selected sequences. Yellow colours indicate higher sound levels. **B**) The ‘sensory evidence’ for the soundscape in A (black line) and other example soundscapes (grey lines). A value of 0.5 indicates ambiguous evidence, a value of 1 that all sequences increased.

Before the actual experiment, we determined participants’ perceptual thresholds using three interleaved 2-down 1-up staircases. An average of six reversals (excluding the initial four) was calculated from each staircase, and the resulting three thresholds were averaged to yield the final participant’s threshold.

The participant’s task was to report the perceived direction of frequency change (‘sweep’) of the stimulus after each trial as accurately as possible. Each experiment consisted of five blocks with 200 trials each, resulting in 1000 trials per participant. The inter-trial intervals had a duration of 1100 ms. Participants could take breaks in between blocks. This design corresponds to Experiment 3 in Kayser et al. 2019, except that here we obtained 1000 trials (rather than 800).The behavioral performance was quantified as the fraction of correct responses and using two measures from signal detection theory: sensitivity (d’) and bias (c)^31^.

### Analysis of psychophysical weights

The soundscapes were designed to allow the application of psychophysical reverse correlation to quantify the influence of the momentary sensory evidence (deviation from an ambiguous sweep direction) on participant’s responses^32,33^.

For each participant we derived a perceptual weighting function as follows: we split trials according to the participants’ response and sweep direction. For each response we calculated the average motion evidence and converted their difference into a within-participant z-score based on a distribution of 4000 weights obtained by randomizing the alignment of stimuli and responses^34,35^. A weight of zero indicates no influence of the stimulus in that epoch on participant’s responses, while positive values indicate a positive relation between sensory evidence and the response. Note that this computation assumes that the time course of the perceptual weights is consistent across trials within a participant, as the reverse correlation assigns a fixed weight to each epoch. To relieve this assumption, the main analysis in this study derived the perceptual weights for subsets of trials that were chosen based on the amplitude or phase of EEG signals in a pre-stimulus period, as described below.

To probe whether these perceptual weights exhibited a systematic rhythmic structure, we proceeded as previously^28^. We first extracted non-rhythmic structure such as an offset, a linear ramp and u/v-shaped time courses fixed to the stimulus duration (the latter modeled as cos(2*pi*t*fexp), with fexp= 1/stimulus duration). These were termed ‘trivial’ components as they do not relate to the specific hypothesis of genuine rhythmic structure at relevant timescales above 1 Hz. To quantify whether a rhythmic component at a frequency above 1 Hz significantly contributes above these trivial components to the perceptual template, we compared regression models featuring only these trivial components with models additionally including a rhythmic component (defined by sine and cosine components of a variable frequency between 1.1 Hz and 4 Hz). For each model we derived its log-evidence obtained from the regression for each participant. Model comparison was based on the group-level log-evidence (assuming that participants contribute independently)^36–38^. In a separate analysis we used a Monte-Carlo approach for model fitting and compared models based on the Watanabe-Akaike information criterion (WAIC), which captures the out-of-sample predictive power when penalizing each model^37^. This calculation was implemented using the Bayesian regression package for Matlab^39^, using 10,000 samples, 10,000 burn-in samples and a thinning factor of 5.

### EEG recordings and analysis

EEG was recorded continuously using a 64-channel ActiveTwo system (Biosemi), with reference electrodes located occipital-parietal at a frequency of 1024 Hz. Electrodes to record the electro-oculograms (EOG) were put below and next to the lateral canthus of both eyes.

The EEG data were analyzed using Matlab (R2017a; TheMathWorks) and the fieldtrip toolbox, version 20190905^40^. The raw data was filtered (between 0.6 and 70 Hz; 3rd-order Butterworth filter) and re-sampled to 150 Hz. Trials were rejected if the amplitude in central electrodes exceeded +/− 175 μV. An average of 21.1 ± 8 (SEM) trials per participant were rejected. Few bad channels were interpolated based on all neighboring channels^41^. Furthermore, artefacts caused by eye movements were identified and rejected, using the data from the EOG channels, and based on an independent component analysis (ICA). Artifacts were identified as in our previous studies^42,43^ following definitions provided in the literature^44,45^. On average we removed 17±2 (mean±s.e.m.) components. Our main analysis focused on the relation between rhythmic brain activity prior to the stimulus and the perceptual weights. To quantify this, we first performed a time-frequency analysis on single trial EEG activity in a time window prior to stimulus onset (−1 s to 0 s). To avoid contamination by post-stimulus activity, we time-mirrored the epoched data and applied a Hanning window to fade out the stimulus period^46^. Time-frequency resolved activity was obtained using Morlet wavelets (4 cycles width) between 2 Hz and 13 Hz, from which we derived the time-varying power and phase of each frequency band. This range was chosen based on the relevant time scales revealed by previous work^8,9,23,47–52^ (or for review see^53^).

### Linking EEG and behavior

To link pre-stimulus EEG activity and behavior, we first quantified the relation between measures of perceptual performance and EEG power and phase. For this, we divided trials into those with high or low pre-stimulus power (based on a median split) for each participant, electrode, frequency and pre-stimulus time point. Then we quantified behavioral performance separately for trials with low or high power. To test for a statistical effect, we then computed a two-sided t-test across participants between low- and high-power bins for each channel, time point and frequency. Because this involved a large number of dimensions, we reduced this dimensionality as follows: we first used the analysis focusing on the fraction of correct responses to define a suitable time point to extract power by finding the time point containing the most significant (at an uncorrected p<0.05) number of channels across frequencies. We then focused on the averaged data power around this time point (−200 ms ±100ms) and tested for a significant relation between power and behavioral sensitivity or bias using cluster-based permutation statistics (see below). To visualize the dependency of behavior on power we also divided the trials according to pre-stimulus power into four bins, with each bin containing the same number of trials.

We used a similar two-stage procedure to test for a relation between EEG phase and behavior. First, we split trials into correct or incorrect responses and contrasted these using the phase opposition sum (POS, VanRullen 2016)^54^. The POS was computed for each participant individually and these were combined across participants using the Stouffer Method using the PhaseOpposition toolbox (VanRullen 2016)^54^. To reduce dimensionality, we calculated the number of significant channels (at p<0.05, uncorrected) for each time point and frequency and determined the time with the highest number of significant channels across frequencies. This time point (−320 ms) was then used for subsequent analyses. For the main analysis we then grouped trials according to phase, similar to the analysis of power. However, because the division of phase in two bins is arbitrary, we repeated this analysis using four different division boundaries to split the circular phase range in two groups (dividing trials along boundaries at 0 and π, along boundaries at π/4 and −3 π/4, etc.). Then we computed the absolute difference in sensitivity or bias between opposing phase bins, selected for each participant, electrode and frequency the phase division yielding the strongest effect, and used cluster-based permutation statistics to determine significant effects.

### Linking EEG and perceptual weights

To test whether and how pre-stimulus power and phase affect the sampling of acoustic information, we quantified the relation between these and the perceptual weights. To do so, we recomputed these weights and their trivial and rhythmic components using regression separately for trials falling into either of the two bins defined based on EEG power or phase. This was done for each participant, frequency band and electrode separately, with phase and power extracted at the respective optimal time points. We then focused on the different components of these weights, considering the rhythmic component at the best group-level frequency of 2.2 Hz (as revealed in **Fig. 2**), and asked whether these components differed in amplitude (or for the rhythmic component, additionally differed in phase) between trials characterized by the two bins of EEG power or phase.

**Figure 2.**
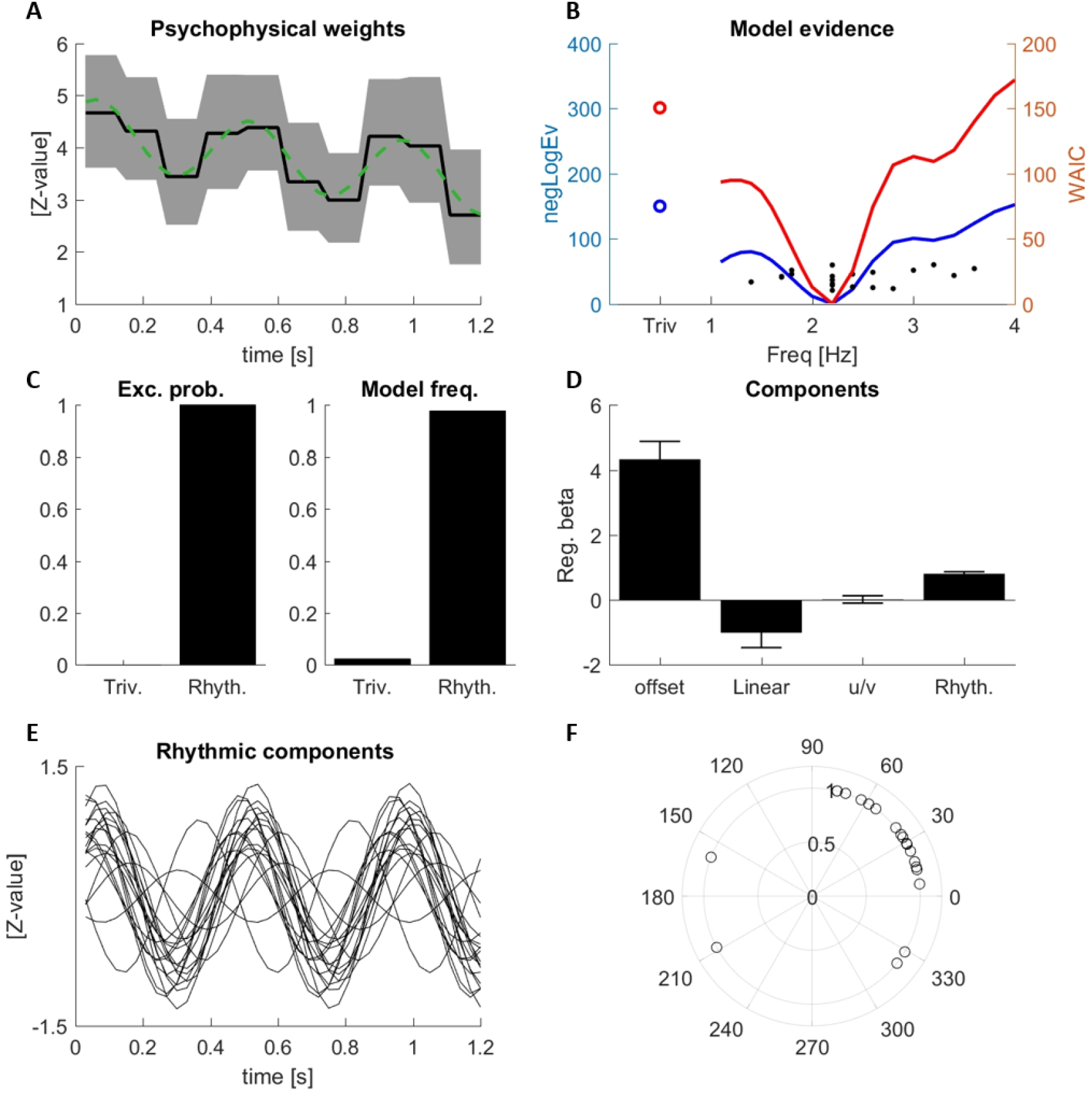
Psychophysical weights. **A)** Group-averaged weights (black line), two-sided 95 % bootstrap confidence interval (grey area) and the best fitting model (green). **B)** Model comparison between the trivial model (circle) and rhythmic models at different frequencies (lines) based on the group-level negative log-evidence (blue) and the WAIC (red). Dots indicate participant’s individual best frequencies (from negLogEv). **C)** Exceedance probabilities and model frequencies comparing the trivial and a rhythmic model at the best rhythmic group-level frequency (2.2 Hz). **D)** Regression betas for each model component (mean, s.e.m.). For the rhythmic component the root-mean-squared amplitude of sine and cosine components at 2.2 Hz is shown. **E)** Rhythmic components of the best model for each individual participant. **F)** Phase of the rhythmic component for each participant.

### Statistics

Statistical tests for EEG data were based on a two-level procedure and used cluster-based permutation procedures to correct for multiple comparisons across electrodes and frequency bands, and for phase, for the inclusion of four potential divisions of phase into two bins. To additionally control for multiple comparisons across performance indices (e.g. d’ and bias; or different components of the perceptual weights) we used the false discovery rate^55,56^ to threshold effects at a corrected p-value of 0.01.

To test for significant relation between EEG power and behavior, we first used a paired t-test to contrast sensitivity (or bias) between power bins across participants. Then, we entered the respective t-values (thresholded at a two-sided level of p<0.05) into a permutation procedure, relying on 2000 permutations of the effect sign across participants, using the max-sum as cluster-forming statistics and considering only clusters exceeding a minimal cluster size of two. The same procedure was used to test the relation between EEG power and parameters derived from the behavioral templates (except the phase of the weighting function; see below).

For EEG phase we used a similar statistical procedure. However, as the split of phase into two bins is arbitrary, we considered for each electrode, frequency and participant four potential divisions of phase into two bins. Because the label of each bin, and hence the sign of the difference of effects between bins is also arbitrary, we computed the absolute difference between phase bins of the variable of interest. We then selected, for each electrode, frequency and participant the one (of four) phase divisions with the largest effect and computed the average across participants. We then compared this true group-level average effect to a distribution of group-level effects obtained from a permutation of trial labels and behavioral data and accepted as significant effects exceeding the 95^th^ percentile of the randomized distribution. We then applied cluster-based permutation procedure as above. The effect of EEG power or phase on the phase of the perceptual weights was tested similarly, by using the absolute value of the change in phase of the weighting function between the two bins derived from EEG power or phase.

## Results

### Behavioral results

Participants were judging the perceived direction of frequency sweep in 1.2 s long soundscapes. These soundscapes **(Fig. 1A**) consisted of 30 simultaneous tone sequences, which varied in frequency, and were designed to allow the quantification of the stimulus-response relation using psychophysical reverse correlation. The resulting perceptual weights are shown in **Figure 2A** and reflect the apparent influence of the momentary sensory evidence on behavior. These weights were significant for all time points (at p<0.05, group-level bootstrap test).

As in our previous study, we investigated the temporal pattern of these perceptual weights by probing their temporal structure using regression modeling. In particular, we asked whether these weights feature rhythmic temporal structure at a time scale of above 1 Hz. To this end, we modeled weights based on three trivial components: a constant offset, a linear slope and a u/v-shaped component time-locked to the sound scape duration, and by adding a rhythmic component with a time scale between 1.1 and 4 Hz. For each participant we quantified the contribution (regression betas) of these four components to the participant-specific weight profile. We then computed the log model evidence for either a regression model comprising only the three trivial components and a model additionally including the rhythmic component at varying frequencies (**Fig. 2 B, blue curve)**. In a separate analysis, we performed the same model comparison using Monte-Carlo simulations used to derive the WAIC criterion (**Fig. 2 B, red curve**). Both analyses consistently revealed that including a rhythmic component at 2.2 Hz provided the highest explanatory power compared to all other frequencies tested. In particular, a group-level model comparison between the trivial model and the rhythmic model at the best group-level frequency (2.2 Hz) revealed that the model including the rhythmic component explained the data better than a model without: the group-level log-evidence was clearly in favor of the rhythmic model (Delta_neglogEv = 147; exceedance probability p_ex_=1; model frequency across participants 0.975; **Fig. 2 C**; the same conclusion was supported by a model comparison based on the WAIC: D_WAIC = 148). This result confirms our previous data that were obtained in a separate group of participants (Experiment 3 in Kayser 2019)^28^.

To illustrate these four contributions to the perceptual weights, the green dashed line in **Figure 2 A** shows the best (group-level) model fit to the actual data. **Figure 2 D** displays the amplitudes (regression beta’s) for the different components: offset (mean= 4.35; SEM= 0.528), linear decrease (mean= −1.002; SEM= 0.461), a small u/v-shaped component (mean =0.026; SEM= 0.117), and a rhythmic component (mean= 0.804; SEM= 0.059). **Figure 2 E** displays the rhythmic component for each participant individually, illustrating the consistency of the rhythmic perceptual weight for most participants. As shown in **Figure 2 F** these rhythmic components share a common phase across most participants.

To confirm whether the behavioral data fit our expectations given the experimental design around participant’s thresholds, we used signal detection theory. Hit rates were around 72% as expected (median= 0.716), sensitivity was above 1 for most observers (Median = 1.173; max= 1.81; min= 0.632) and the response criterion revealed no bias (median= −0.01; max= 0.299; min= −0.619).

### EEG results

The analysis of EEG data was designed to probe whether these perceptual weights reflecting the influence of acoustic evidence on participants’ behavior were related to the state of rhythmic brain activity prior to the stimulus. That is, we asked whether (statistically) the shape of these weights differed depending on the state of brain activity prior to the stimulus. Such a dependency could for example reveal whether the overall strength of perceptual sampling (offset), or the strength of the rhythmic contribution differs between trials with particularly high or low power. It could also reveal whether the timing of rhythmic sampling (the weight’s phase) differs between trials preceded by a particular phase in a specific EEG band.

Addressing these questions using statistics required us to first reduce the complexity of this analysis by removing one (least-interesting) dimension: the precise time point prior to the stimulus used to characterize the brain activity. We hence implemented a first analysis determining the time points that seemed most promising to capture any dependency between EEG power (or phase) and behavior. To do so for EEG power we computed the difference in the fraction of correct responses between trials with high or low power and quantified how many channels exhibited a significant difference (at p<0.05) at each frequency and time point. This revealed a cluster of more than 60% significant channels between 300 ms to 100 ms before stimulus onset, with a peak around 200 ms. We hence used the average over this time window for the subsequent analysis of EEG power. For EEG phase, we calculated the phase opposition sum (POS) as an index of whether pre-stimulus phase differs between trials with correct and wrong responses^54^. Calculating the number of channels with a significant group-level effect revealed a peak 320 ms before stimulus onset, which was used for the subsequent analysis of EEG phase.

### Linking EEG activity and behavior

To test for a relation between EEG power and behavior, we contrasted perceptual sensitivity and bias between bins of high and low power (**Fig. 3 A-E**). For sensitivity this revealed a significant positive cluster between 8 and 13 Hz (t_clus_=449.016 p=0.002, 42 channels over fronto-central areas), which was also significant after correcting for all comparisons using the False Discovery Rate (p<0.01). The distribution of this cluster is visualized by highlighting all significant electrodes in one topographic plot (**Fig. 3 C**). For bias we did not find a significant effect (**Fig. 3 B**). To visualize the relation between EEG power and behavior in more detail, **Figure 3 D, E** show the sensitivity/bias as a function of power, obtained by dividing trials according to power into four bins.

**Figure 3.**
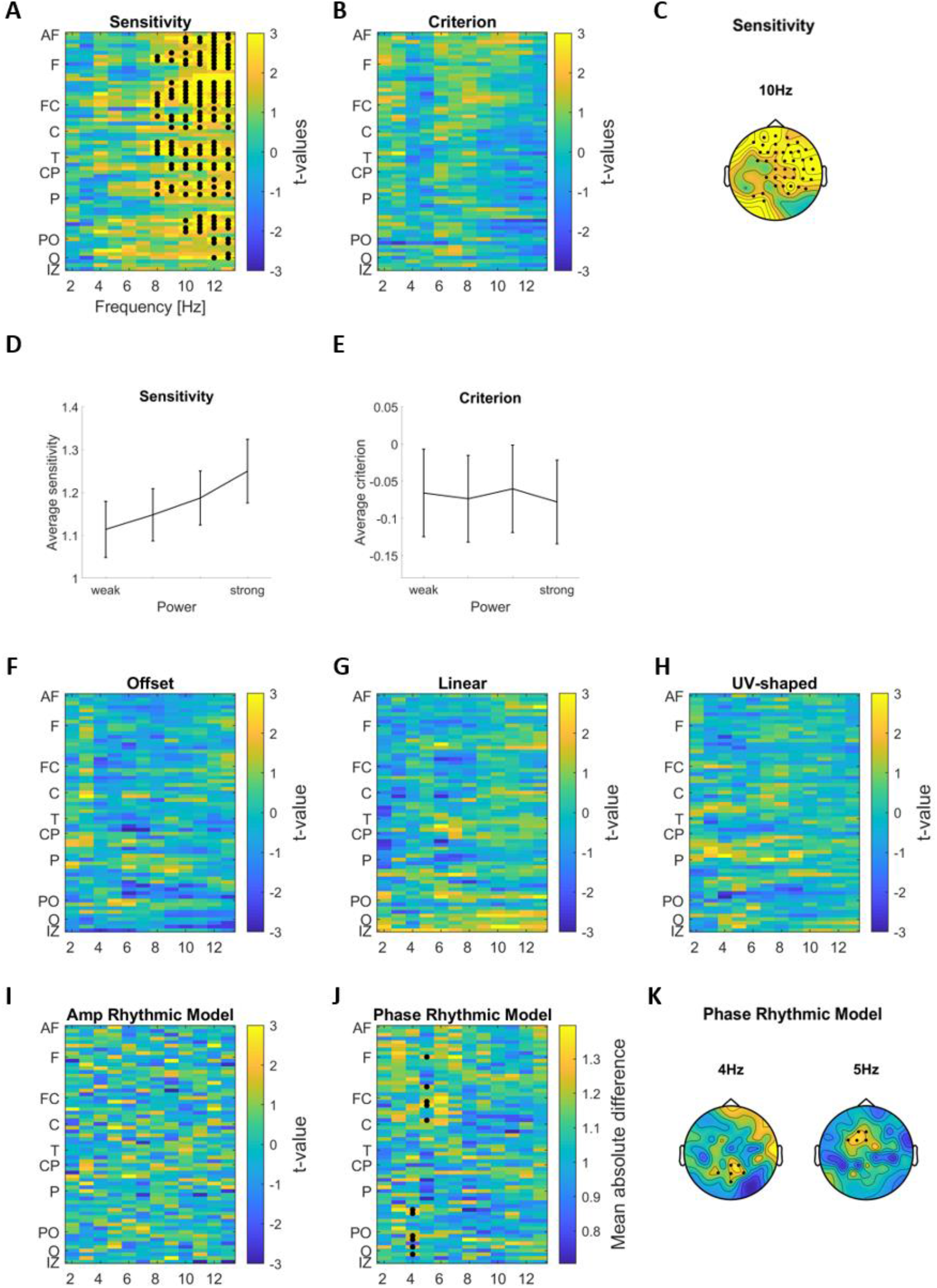
Linking EEG power with behaviour and psychophysical weights. **A-B)** Difference in perceptual sensitivity and criterion between trials with high or low pre-stimulus power (group-level t-values; paired t-test). **C)** Topography of the significant electrodes for sensitivity overlaid on the t-map at 10Hz. **D, E)** Sensitivity and criterion across participants and significant channels (from the cluster for sensitivity in panel A; mean and s.e.m. across participants) as a function of power (four equi-populated bins). **F-I)** Difference in the prominence (amplitude) of the different components of the perceptual weights between trials with high/low pre-stimulus power (group-level t-values; paired t-test), as a function of EEG band and EEG electrode. **J, K)** Difference in relative phase of the rhythmic sampling component between trials with high/low power (group-level average absolute difference). Electrodes from significant clusters are marked with black dots (first level significance at p < 0.05, cluster significance at p < 0.01 FDR).

We performed a similar analysis for EEG phase (**Fig. 4 A-B**). This revealed no significant effects (at p<0.01).

**Figure 4.**
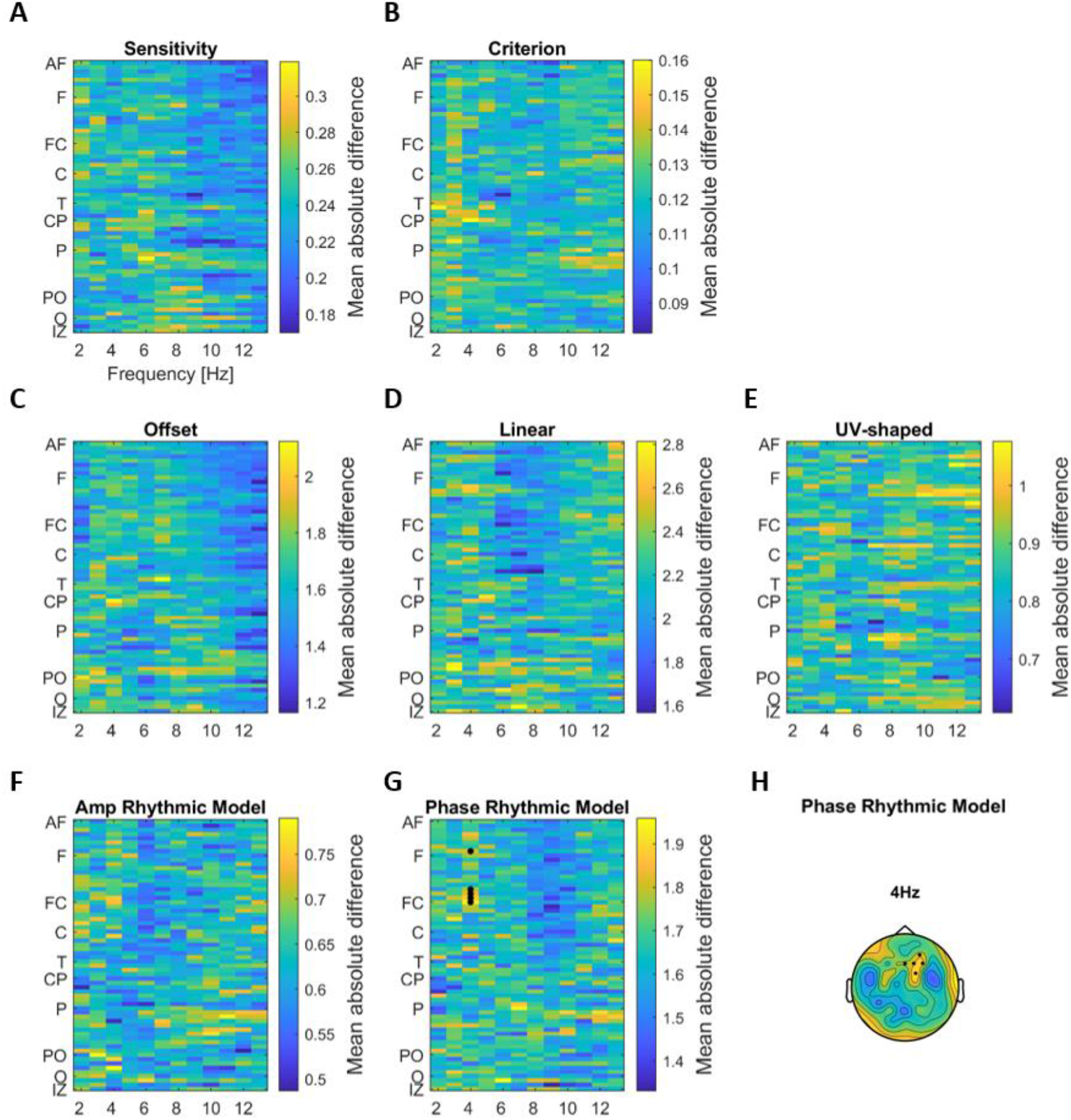
Linking EEG phase with behavior and perceptual weights. **A, B)** Group-level average of participants-wise differences in perceptual sensitivity and criterion between trials with opposing EEG phase. **C-G)** Group-level average of participants-wise differences in regression betas of each component of the perceptual weights between trials with opposing EEG phase, as a function of EEG band and EEG electrode. **H)** Topographic representation of the group-level average of participants-wise differences in the phase of the rhythmic model. Electrodes from significant clusters are marked with black dots (first level significance at p < 0.05, cluster significance at p < 0.01 FDR).

### Linking EEG activity and perceptual weights

To test whether the perceptual sampling of acoustic information is affected by pre-stimulus activity, we asked whether the psychophysical weights differ between trials characterized by high or low EEG power, or by different phase bins in a particular frequency band. We tested such relations for each of the four model components used to describe the weighting function (offset, linear slope, u/v curve, and rhythmic component). In doing so, we focused on the amplitude of all components and for the rhythmic component also on the relative phase angle of this. The latter analysis allowed us to directly test whether for example the pre-stimulus phase affects the phase of the rhythmic perceptual sampling. Statistical cluster-based permutation tests were corrected for multiple frequencies and considering multiple division of phase into two bins using the max-statistics and for multiple contrasts using the FDR.

For pre-stimulus power we found no significant effects for the trivial model parameters and the amplitude of the rhythmic model (**Fig. 3 F-I**). However, the tests revealed a significant effect for the phase of the rhythmic model in the 4 and 5 Hz EEG bands (t_clus_=11.787 p=0.0017, 6 channels over parieto-occipetal areas; t_clus_=10.473 p=0.005, 5 channels over fronto-central areas), which were also significant after correcting for all (p<0.01) (**Fig. 3 J, K**).

For phase, these tests revealed no significant effects on the three trivial model parameters (offset, linear and u/v-shaped) and the amplitude of the rhythmic model (at p<0.01) (**Fig. 4 C-F**). However, a significant cluster emerged for the influence of the relative phase of the rhythmic component in a frontal cluster at 4 Hz (**Fig. 4 G, H**; t_clus_t= 11.998, p<0.001, 5 channels over audio-frontal and frontal areas). This indicates that the relative phase of how perception samples acoustic information around 2 Hz changes with the relative phase of the EEG activity over frontal regions at about 4 Hz. Given that both EEG phase and power at 4Hz revealed a significant relation to the phase of rhythmic perceptual sampling, we asked whether the strength of both effects was correlated across participants. A non-parametric correlation turned out to be not significant (rank-correlation: r= 0.4331, p=0.0565, 95^th^ bootstrap-CI [-0.0144, 0.7097]).

To visualize the relation between pre-stimulus EEG power/phase and the perceptual sampling weights we reconstructed the rhythmic component of these weights for the participant’s specific EEG bins individually. Four examples are shown in **Fig. 5 A, B**, illustrating both the different sampling phase between EEG-power/phase bins, as well as the heterogeneity of this effect across participants. On average across participants the absolute phase shift in rhythmic sampling across opposing EEG-power bins was 75.68° (95^th^ CI of the mean [57.19, 94.15]) (**Fig. 5C**). The absolute phase shift in rhythmic sampling across opposing EEG-phase bins was 110.98° (95^th^ CI of the mean [92.33, 129.64]) (**Fig. 5D**).

**Figure 5.**
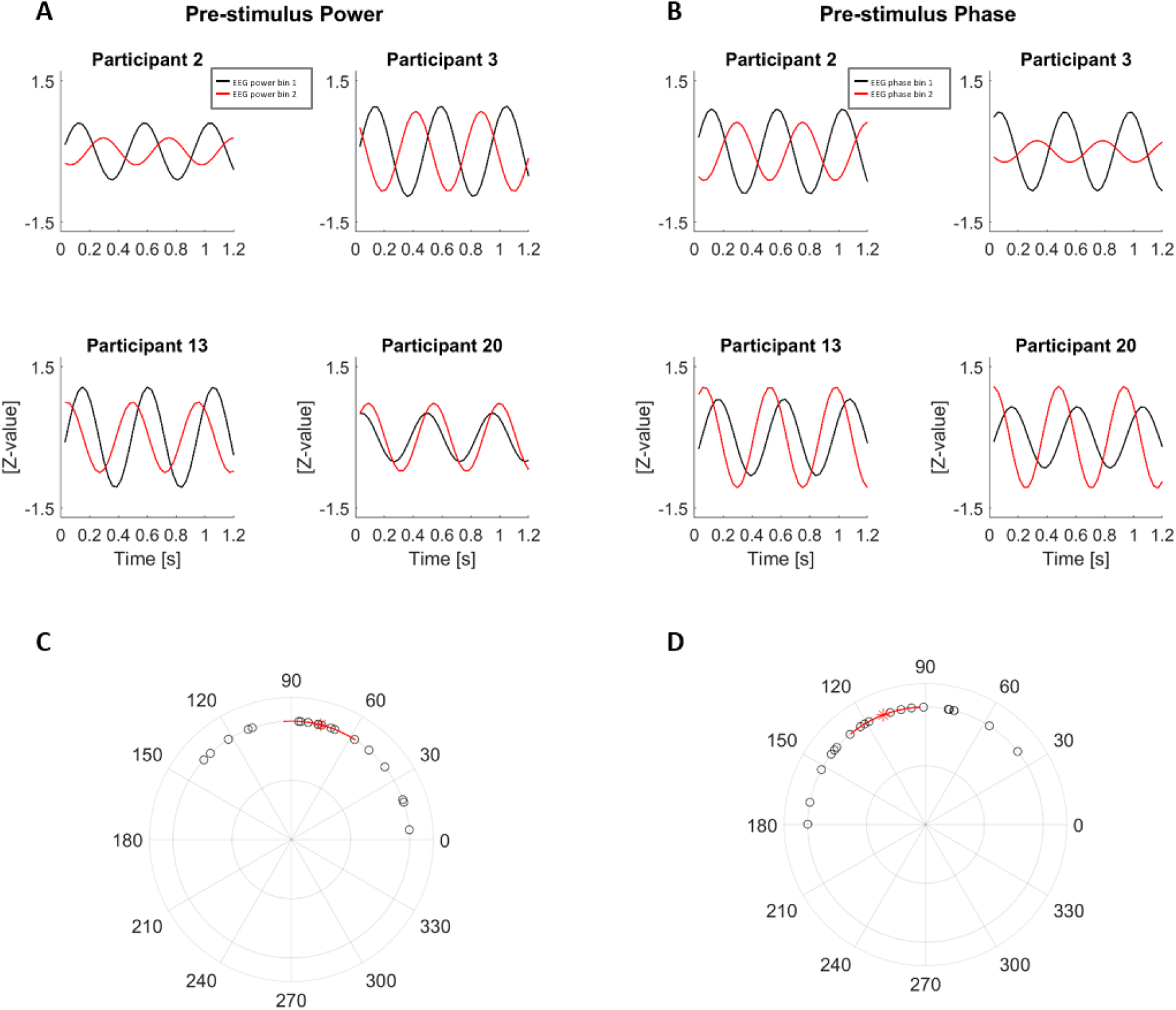
Visualization of the change in rhythmic sampling with EEG power/phase. **A, B**) Examples of the rhythmic components (at 2.2 Hz perceptual sampling frequency) of the perceptual weights for four participants reconstructed for the respective participants-selected EEG-power **(A)** and phase **(B)** bin. For this we considered the EEG phase bins at 4 Hz of each participant yielding the largest difference, derived from the electrode in the significant cluster exhibiting the strongest (Pz (power) /FC2 (phase)). **C, D**) Absolute phase differences in the rhythmic perceptual weights between the two EEG-power **(C)** /phase **(D)** bins for each participant. The group-mean is marked with the red star and 95% confidence intervals (CI) are marked by the red lines.

## Discussion

We investigated whether brain activity prior to a stimulus influences the manner in which human participants use the moment by moment acoustic evidence to make a perceptual judgement pertaining to temporally extended auditory scenes. Confirming previous results, we found that participants sample the acoustic evidence not uniformly^28^; rather, the weights characterizing the perceptual sampling reveal a rhythmic pattern at a frequency of about 2 Hz. Importantly, the phase of this perceptual sampling co-varied with the state of pre-stimulus brain activity around 4 Hz, suggesting that the rhythmic sampling process observed in the behavioral data is directly linked to ongoing (rhythmic) brain activity. We also found that perceptual performance varied with the strength of pre-stimulus alpha band power over frontal electrodes, demonstrating a general influence of pre-stimulus activity on perception for prolonged stimuli. These results support the notion that the state of delta/theta band brain activity shapes the manner by which subsequent acoustic evidence influences auditory perception.

### Evidence for the Rhythmic Sampling of Auditory Scenes

The present study capitalized on a previously developed paradigm and the present data reproduce our previous results^28^. In particular, we show that the perceptual weights attributed to different epochs in temporally extended soundscapes (1.2 s here) exhibit a rich temporal structure, including but not limited to, a linear trend and a significant rhythmic component at a group-level frequency of 2.2 Hz. This conclusion is supported by two different approaches to determining whether including any rhythmic structure better explains the perceptual weights than models not containing such rhythmic structure. For the same soundscape duration (Experiment 3 in Kayser et al 2019)^28^ the previous study revealed a very similar sampling frequency of 2 Hz. Importantly, the acoustic soundscapes used in this experiment are devoid of specific rhythmic structure at this time scale^28^, suggesting that the apparent rhythmic perceptual sampling must be driven by some endogenous mechanism operating at the delta band time scale.

However, the analysis of the behavioral data necessitates the assumption that the perceptual weight attributed to each epoch is fixed across trials, because the perceptual weight is estimated by combining the sensory information and behavioral outcome across trials. However, this assumption may not be valid, in particular if the hypothesized link between delta/theta band brain activity and rhythmic modes of perception is true. To overcome this limitation, we combined single trial estimates of pre-stimulus brain activity with the psychophysical reverse correlation estimate.

### Pre-stimulus activity shapes the timing of perceptual sampling

Our results directly reveal a correlation between the power and phase of pre-stimulus delta/theta band activity around 4 Hz and the relative timing of the subsequent perceptual weighting process. Thereby our results provide a direct link between the pre-stimulus brain state and the subsequent exploitation of acoustic information for active listening over prolonged epochs. At the same time, we note that the strength of the phase-shift in perceptual sampling across opposing power/phase bins of the EEG activity was variable across participants (ranging from 5 to 138 degrees for power and from 39 to 180 degrees for phase). This suggests that the underlying effect is either highly variable across participants or that the effect sizes obtained in the present study are still limited by number of trials collected.

Previous work has linked auditory delta band activity with changes in both spontaneous and stimulus driven neural activity^20,29,57^. The engagement of delta band activity has been implied in the attentional filtering of soundscapes and the task-relevant chunking of speech sounds into sentence or word-level structures^58–60^, and plays a central role in theories of rhythmic modes of listening^2,17,19^. However, a critical hypothesis emerging from these studies had not been tested: that rhythmic pre-stimulus activity is directly linked to the subsequent perceptual use of acoustic information over prolonged time scales. While some studies have shown the persistent fluctuations of behavioral performance at similar time scales subsequent to a brief stimulus, no study has shown that pre-stimulus activity shapes how temporally extended soundscapes are sampled to make a perceptual decision. We here close this gap by directly linking pre-stimulus activity and the subsequent perceptual influence of this.

### Pre-stimulus power shapes behavior

We also found a significant relation between pre-stimulus alpha (8-13 Hz) power and participants’ sensitivity to the direction of acoustic sweep. Generally, such a relation of pre-stimulus power and perceptual performance has been observed in a wide range of perceptual studies across sensory modalities^8,14,61–66^. However, most studies in the auditory domain reported effects predominantly at lower delta or theta band frequencies^8,9,11–14^, while a role for alpha band activity is typically discussed for vision and spatial attention paradigms.

Previous work on the entrainment of brain activity to speech has shown that frontal alpha band power correlates with the strength of delta band speech-tracking^20^. Frontal alpha could reflect a mechanism that shapes the alignment of rhythmic activity in auditory regions to the acoustic stimulus in a top-down manner^67,68^. Given that stronger speech-to-brain alignment is also predictive of an improved speech reception^69,70^ these results suggest that frontal alpha band activity may be more generally predictive of the correct identification of complex sounds. The positive relation of pre-stimulus alpha and improved sensitivity observed here provides further support for this, although we did not find a significant relation between alpha power and the perceptual weights themselves.

Alternatively, these discrepancies in perceptually-relevant EEG frequencies in the present and previous work could be explained by the rather long duration of the soundscapes used here. While the stimuli here lasted more than a second, previous studies reporting a correlation of pre-stimulus power and behavioral outcome mostly used very brief stimuli (e.g. < 200ms)^8,14^. Hence, one cannot rule out that the previously observed effects in delta/theta bands and the alpha effect shown here reflect two different neurophysiological mechanisms that each shape perception for shorter and longer stimuli, respectively.

## Conclusion

We systematically investigated the relation between pre-stimulus brain activity and rhythmic perceptual sampling of long and non-rhythmic stimuli. Our data show that strength and the timing of delta/theta band pre-stimulus EEG activity relates to the rhythmic perceptual sampling of auditory scenes. These results directly point to a lasting influence of spontaneous rhythmic brain activity for the perception of subsequent stimuli and close a critical gap in the mechanistic picture proposing a fundamental role of rhythmic auditory cortical activity for active listening.

## Acknowledgements

C Kayser is supported by the European Research Council (to C.K. ERC-2014-CoG; grant No 646657).

## Author contributions

C. Kayser designed the experiment, C. Kubetschek collected the data, both authors analysed the data and wrote the manuscript

## Additional Information

Competing interests: The authors declare no competing interests.

## Data availability

The original and preprocessed data and Matlab code are available upon request.

## Notes

### Competing Interest Statement

The authors have declared no competing interest.

